# Specific Aneuploidies Predict Immune Evasion and Poor Immunotherapy Response in Melanoma

**DOI:** 10.64898/2026.04.12.718039

**Authors:** Lizabeth Katsnelson, Shini Chen, Ze Chen, Dania Annuar, Jesus Mario Rangel Valenzuela, Andy Zhao, Joanne Xiu, David Fenyő, Joy Bianchi, Teresa Davoli

## Abstract

Melanoma is one of the leading cancer types treated with immune checkpoint blockade (ICB), yet a substantial proportion of patients fail to respond. While tumor mutational burden and PD-L1 expression are established ICB biomarkers, they leave large gaps in predictive accuracy. Somatic copy number alterations (SCNAs) are pervasive in melanoma but their role in shaping the tumor immune microenvironment (TME) and predicting immunotherapy outcomes has been insufficiently characterized.

Here we present KaryoTME, an integrated computational framework that systematically links SCNAs to immune phenotypes using genomic, transcriptomic, and clinical data from over 15,000 patients. Applying this framework to skin melanoma (SKCM) within a pan-cancer context, we identify arm-level chromosome 1q gain and 9p loss as the most prominent SCNA events associated with an immune-cold tumor microenvironment. These alterations act through distinct mechanisms: 9p loss preferentially depletes NK and CD8^+^ T cells, whereas 1q gain is more strongly associated with reduced anti-tumor immune cell infiltration. At the focal level, regions 1q21 and 1q42 show the strongest immune-suppressive associations in melanoma.

Applying the TUSON-Immune algorithm, we predict candidate Tumor immune Suppressor Genes (TiSG) and immune Oncogenes (iOG) within these chromosomal regions, revealing enrichment for pathways including IFN signaling, JAK/STAT pathway, and immune-suppressive cytokine secretion.

Critically, 1q gain emerged as a strong and independent predictor of poor survival following anti-PD-1/PD-L1 therapy across two independent clinical cohorts: the MSK-IMPACT cohort (p = 0.018, N = 77) and a large real-world Caris Life Sciences dataset (HR = 1.2, p = 0.002, N = 1,167). Multivariate analysis confirmed that 1q gain predicts poor outcomes independently of CD8^+^ T-cell infiltration, B-cell infiltration, tumor mutational burden, and PD-L1 status. These findings establish chromosome 1q gain as a compelling biomarker of immunotherapy resistance in melanoma and highlight aneuploidies as underappreciated drivers of immune evasion in this disease.

## Introduction

Melanoma is among the most immunogenic solid tumors, and the development of immune checkpoint blockade (ICB) therapies targeting PD-1/PD-L1 has transformed its clinical management. Nevertheless, only 30–40% of patients achieve durable responses, and the mechanisms underlying primary and acquired resistance remain incompletely understood. While tumor mutational burden (TMB) and PD-L1 protein expression provide some predictive value, they leave substantial gaps, particularly in patients with microsatellite-stable tumors and low mutational loads.

Somatic copy number alterations (SCNAs), which include gains and losses of chromosomal segments ranging from focal amplifications to whole-arm events, represent one of the most pervasive forms of genomic instability in melanoma. Cutaneous melanoma harbors a high burden of SCNAs, with recurrent events including 9p loss (encompassing *CDKN2A, JAK2*, and interferon genes), 1q gain, and 6p gain. While individual loci such as 9p24 (containing *PD-L1*/*PD-L2* and *JAK2*) have been studied in the context of immunotherapy response, a genome-wide, systematic analysis of SCNAs as drivers of immune evasion and predictors of ICB outcomes in melanoma is lacking.

Importantly, SCNAs can affect immune cell infiltration and function through multiple mechanisms. General chromosomal instability (CIN) promotes an immunosuppressive microenvironment via chronic cGAS/STING activation and unfolded protein responses. In addition, chromosome-specific SCNA effects, arising from the combined dosage changes of multiple immune-relevant genes within a chromosomal region, can selectively deplete cytotoxic immune populations or promote immunosuppressive myeloid cells. Disentangling these chromosome-specific effects from global aneuploidy burden is essential for identifying actionable biomarkers.

Here, we developed KaryoTME (Karyotype-driven modulation of the Tumor Microenvironment), an integrated analysis framework that combines SCNA profiling, immune deconvolution, and clinical ICB outcome data across thousands of patients. Using this approach within a pan-cancer context that highlights melanoma, we identify chromosome 1q gain and 9p loss as key SCNA drivers of immune evasion. We characterize the specific immune cell populations affected by these alterations, predict candidate driver genes within these regions using the TUSON-Immune algorithm, and demonstrate that chromosome 1q gain is an independent predictor of poor survival in two large melanoma cohorts treated with ICB therapy. These findings provide both mechanistic insights into SCNA-driven immune evasion in melanoma and a potential biomarker for improving patient stratification.

## Results

### KaryoTME framework identifies SCNA events associated with immune-cold and immune-hot tumor microenvironments

To systematically examine the relationship between chromosomal copy number events and immune infiltration, we developed KaryoTME (Figure 1A–B). For each tumor sample, we computed a cytotoxic Immune Score (IS) based on the ranked expression of seven cytotoxic genes (CD247, CD2, CD3E, GZMH, NKG7, PRF1, GZMK), which collectively capture the extent of CD8^+^ T-cell and NK-cell infiltration (Figure 1B). We then employed logistic regression models to determine, for each tumor type and chromosomal locus, whether a given SCNA (gain or loss) is associated with a low (“immune cold”) or high (“immune hot”) IS, while including the overall aneuploidy score (AS) as a covariate to control for global chromosomal instability. Gain and loss models were run independently to avoid confounding between the two directions of copy number change.

**Figure 1.**
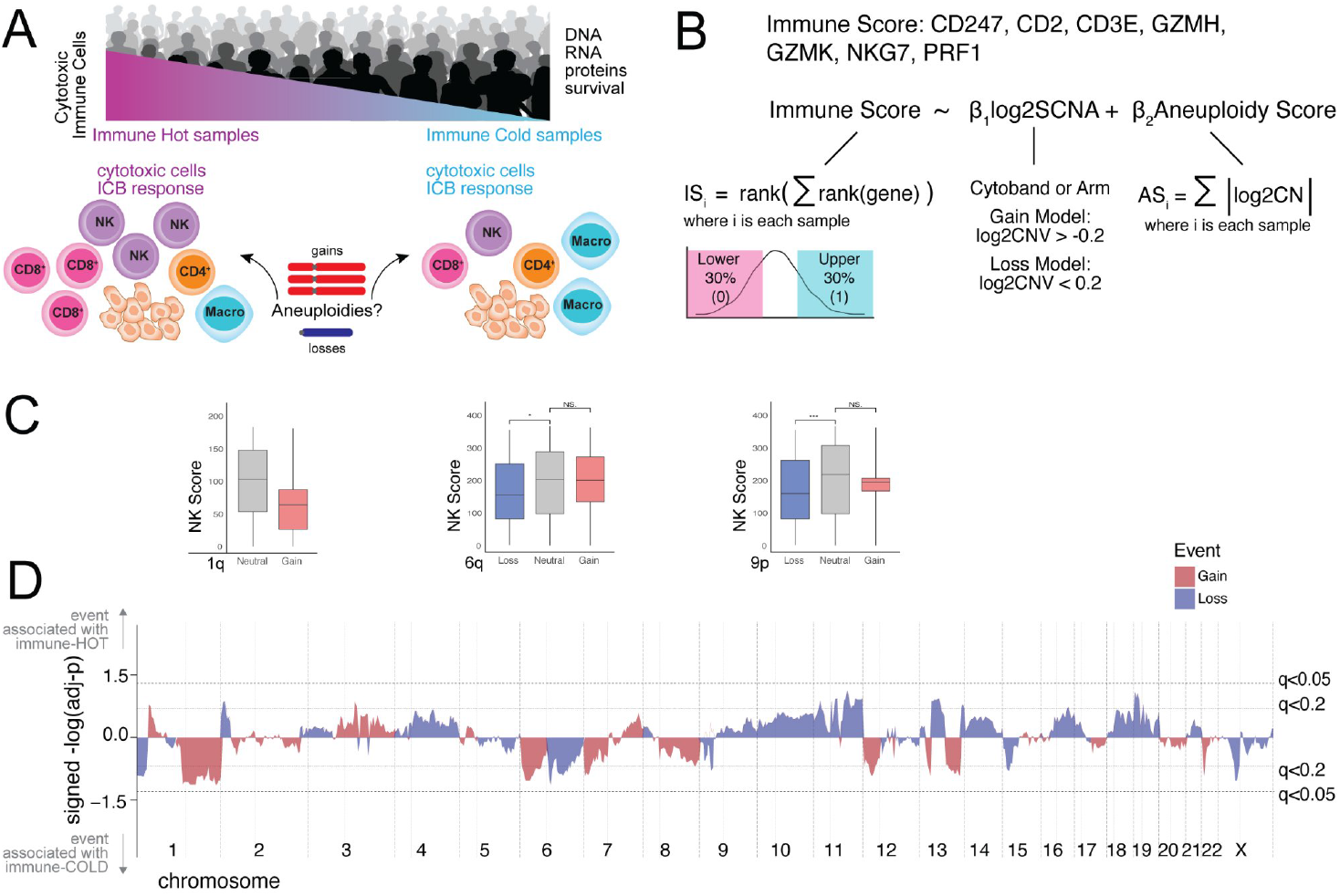
Overview of the KaryoTME framework and SCNA–immune associations in skin melanoma. (A) Schematic of immune hot and immune cold tumor microenvironments and study design. (B) Definition of the cytotoxic Immune Score (IS), the SCNA regression model, and the aneuploidy score (AS). (C) NK-cell score in skin melanoma (SKCM) across SCNA status groups for 1q gain, 6q alterations, and 9p alterations. Boxplots show median and interquartile range; asterisks indicate statistical significance. (D) Genome-wide landscape of SCNA–immune associations in SKCM (signed −log_10_(adjusted-p)). Red bars indicate gains; blue bars indicate losses. Dotted lines mark q < 0.05 and q < 0.2 thresholds. Events above the x-axis are associated with immune-hot TME; events below are associated with immune-cold TME.

As illustrative examples of SCNAs with strong immune associations in SKCM, we examined NK-cell scores in relation to 1q gain, 6q alterations, and 9p alterations (Figure 1C). Consistent with pan-cancer trends, 1q gain showed reduced NK-cell infiltration, and 9p loss was significantly associated with lower NK-cell scores. These relationships highlight the direction-dependent nature of SCNA–immune associations: gains and losses of the same chromosomal region can have distinct effects on immune infiltration.

Figure 1D displays the genome-wide landscape of cytoband-level SCNA–immune associations in SKCM (signed −log_10_(adjusted-p) values). Events above the upper dotted line (q < 0.05) are associated with an immune-hot phenotype, while events below the lower dotted line are associated with immune-cold phenotype. This landscape underscores that 1q gain (immune-cold) and 9p loss (immune-cold) are among the most significant events in melanoma.

### Arm- and focal-level SCNA associations reveal recurrent immune-suppressive events including 1q gain and 9p loss

To characterize the pan-cancer landscape of SCNA–immune associations and place melanoma findings in context, we examined arm-level and focal (cytoband-level) SCNAs associated with immune-cold and immune-hot TMEs across 31 tumor subtypes (Figure 2A–B). Despite widespread tissue specificity, several events were recurrent across multiple tumor types. Among SCNAs associated with immune-cold TME in three or more tumor types, arm-level losses (including 5q, 9p) and gains (including 1q, 6p) were prominent. 9p loss was the single most broadly significant immune-cold event, consistent with its role in removing immune-activating genes including the interferon gene cluster, JAK2, and PD-L1/PD-L2. Chromosome 1q gain was associated with immune-cold TME in lung adenocarcinoma, endometrial cancer, and skin melanoma, among others.

**Figure 2.**
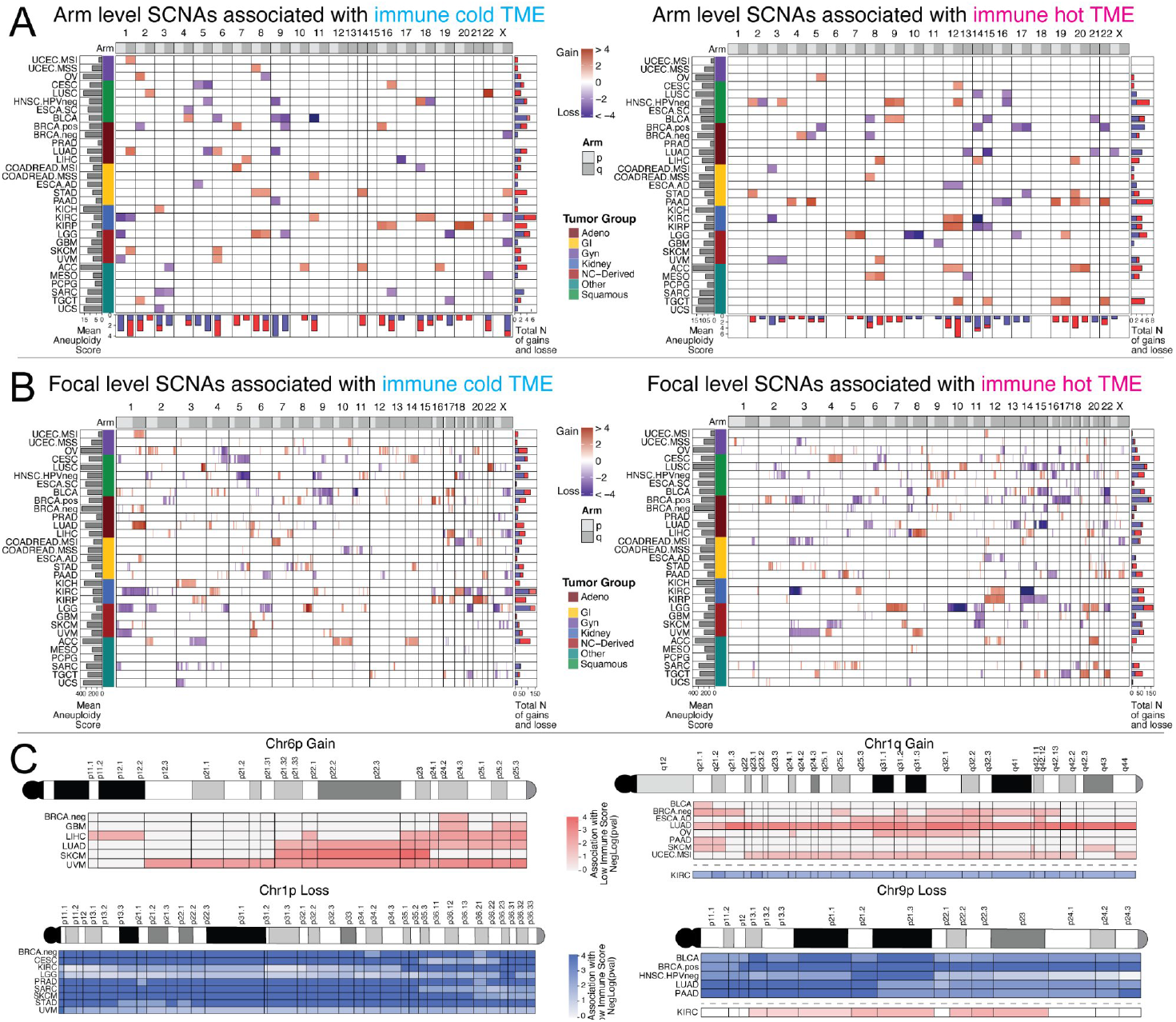
Arm- and focal-level SCNA associations with immune TME across tumor types. (A) Heatmaps of arm-level SCNAs associated with immune-cold (left) and immune-hot (right) TME across 31 tumor subtypes. Color intensity reflects effect size (log2 scale); border shading distinguishes p < 0.05 (solid) from q < 0.05 (bold). Bar graphs at bottom and right show total number of significant events per arm and per tumor type, respectively. (B) Focal (cytoband-level) SCNA associations with immune-cold (left) and immune-hot (right) TME, displayed as in panel A. Cytoband-resolution maps for Chr6p Gain, Chr1q Gain, Chr1p Loss, and Chr9p Loss across selected tumor types. Rows are tumor subtypes; columns are cytobands. Color scale indicates the strength of association with immune-cold TME (−log(p)); pink/red is used for gain events and blue for loss events. Deeper color indicates stronger association.

In skin melanoma specifically, arm-level analysis identified 1q gain, 9p loss, and 6p gain as the top immune-cold events, consistent with their high recurrence in melanoma genomics. At the focal level, two regions of chromosome 1q showed particularly strong associations: cytobands 1q21 (near the centromere) and 1q42 (close to the telomere) were associated with reduced IS in skin melanoma as well as in bladder, breast, and pancreatic cancers (Figure 2C). For chromosome 9p loss, cytobands 9p21, 9p23, and 9p24 showed the strongest associations with immune-cold TME across the tumor types highlighted in Figure 2C, including bladder, breast, head and neck, and lung adenocarcinoma.

Importantly, kidney tumors displayed a striking inversion of the typical SCNA–immune relationships, with 1q gain and 9p gain associated with immune-hot rather than immune-cold TME (Figure 2A, C). This tissue-specific reversal likely reflects the unique immunobiology of renal cell carcinoma and serves as an internal control demonstrating the specificity of our models. Altogether, these data highlight 1q gain and 9p loss as pan-cancer immune-evasion events with particularly pronounced effects in melanoma.

### 1q gain and 9p loss affect distinct immune cell populations, with NK and CD8^+^ T-cell depletion in melanoma

To determine which immune cell populations are most strongly affected by SCNAs, we extended the KaryoTME regression framework to model individual cell-type abundances estimated by xCell deconvolution of bulk RNA-seq data. For each arm- and focal-level SCNA, we quantified the number of significant associations with low or high abundance of CD8^+^ T cells, CD4^+^ T cells, B cells, NK cells, anti-tumor macrophages, pro-tumor macrophages, dendritic cells, neutrophils, fibroblasts, and endothelial cells (Figure 3A).

**Figure 3.**
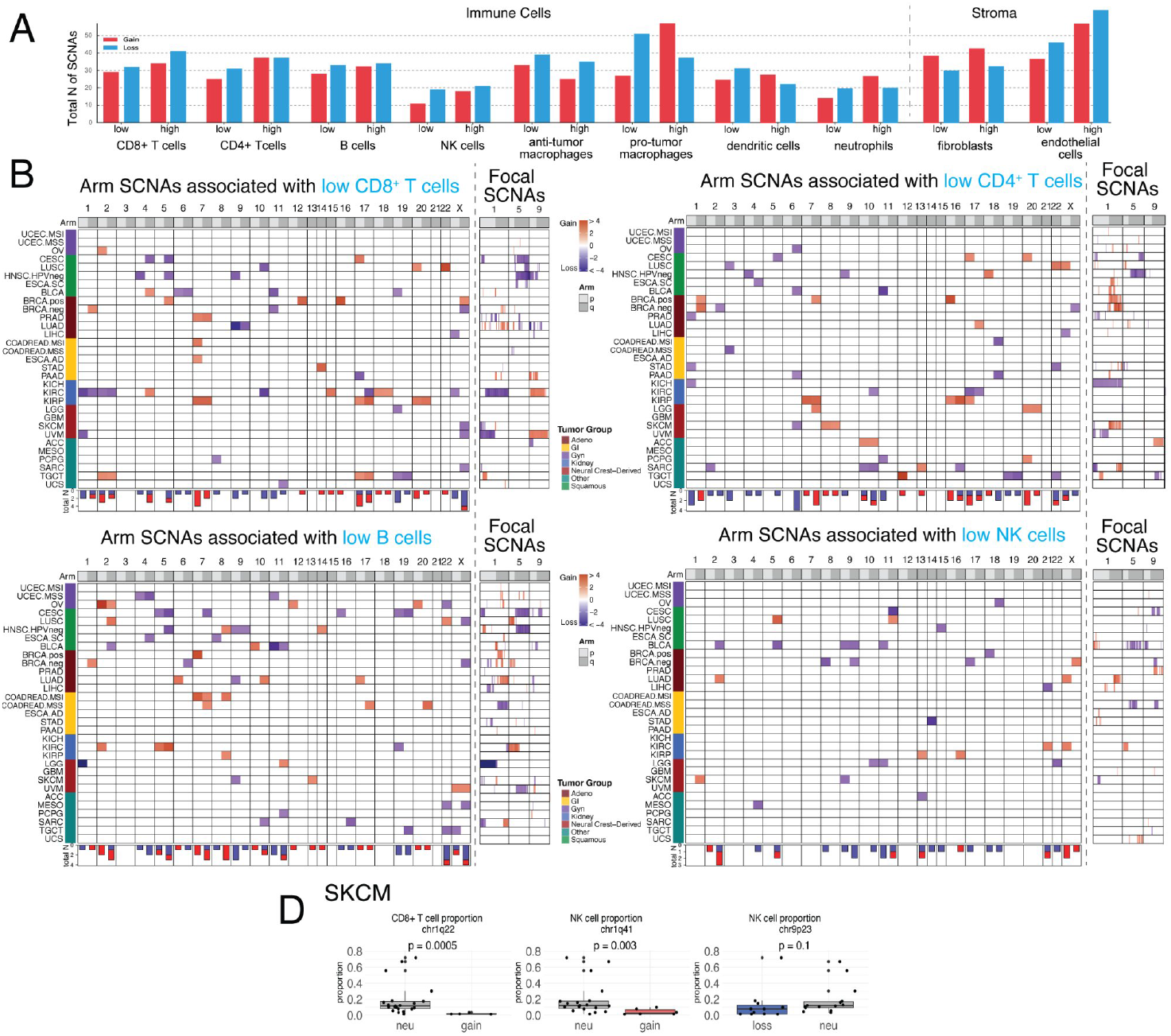
SCNA associations with specific immune cell populations and single-cell validation in melanoma. (A) Total number of significant arm-level SCNA associations with low or high abundance of each indicated immune or stromal cell type across all tumor types. Bar colors indicate SCNA type: Gain (red) and Loss (blue). (B) Heatmaps of arm-level SCNAs associated with low CD8^+^ T cells, CD4^+^ T cells, B cells, and NK cells across 31 tumor subtypes (same layout as Figure 2A). Focal SCNA panels on the right show associations at chromosomes 1, 5, and 9.Single-cell RNA-seq validation in skin melanoma (SKCM). Boxplots show cell-type proportions stratified by SCNA status for chr1q22 (CD8^+^ T-cell proportion; p = 0.0005), chr1q41 (NK-cell proportion; p = 0.003), and chr9p23 (NK-cell proportion; p = 0.1). Each dot represents a patient sample.

Across tumor types, macrophages, both anti-tumor (M1-like) and pro-tumor (M2-like), displayed the highest total number of significant SCNA associations (Figure 3A). T cells showed an intermediate number of associations, while NK cells and neutrophils had relatively fewer. Pan-cancer heatmaps for arm-level SCNAs associated with low CD8^+^ T cells, CD4^+^ T cells, B cells, and NK cells are shown in Figure 3B. Importantly, different SCNA events were preferentially associated with depletion of different immune compartments: 9p loss was predominantly linked to reduced CD8^+^ T cells, B cells, and NK cells (Figure 3B), consistent with direct suppression of cytotoxic immunity. The pan-cancer analysis reveals macrophages as the cell type with the highest total number of SCNA associations (Figure 3A), with gains showing particularly elevated counts for low anti-tumor macrophage abundance across tumor types.

We validated these associations in melanoma using single-cell RNA-seq data (Figure 3D). In SKCM, chromosome 1q22 gain was significantly associated with lower CD8^+^ T-cell proportions (p = 0.0005), and 1q41 gain was associated with reduced NK-cell proportions (p = 0.003). The corresponding effect for 9p23 loss trended in the expected direction but did not reach significance in this relatively small single-cell dataset (p = 0.1). These single-cell analyses provide orthogonal, higher-resolution evidence for the immune-modulatory effects of key SCNAs in melanoma, and confirm that 1q gain affects both the adaptive and innate immune compartments in this tumor type.

### TUSON-Immune predicts candidate Tumor immune Suppressor Genes and immune Oncogenes within melanoma-relevant chromosomal regions

To nominate individual genes driving immune evasion within SCNA-affected chromosomal regions, we applied TUSON-Immune (Tumor Suppressor and Oncogene Navigator–Immune), an extension of the TUSON framework (Figure 4A). Each gene was evaluated using three parameters selected by Lasso classification: the correlation between SCNA and IS (DNA:IS), the correlation between RNA expression and IS (RNA:IS), and the correlation between SCNA and RNA expression (DNA:RNA). Genes in chromosomal losses meeting criteria for Tumor immune Suppressor Genes (TiSGs) and genes in gains meeting criteria for immune Oncogenes (iOGs) were retained.

**Figure 4.**
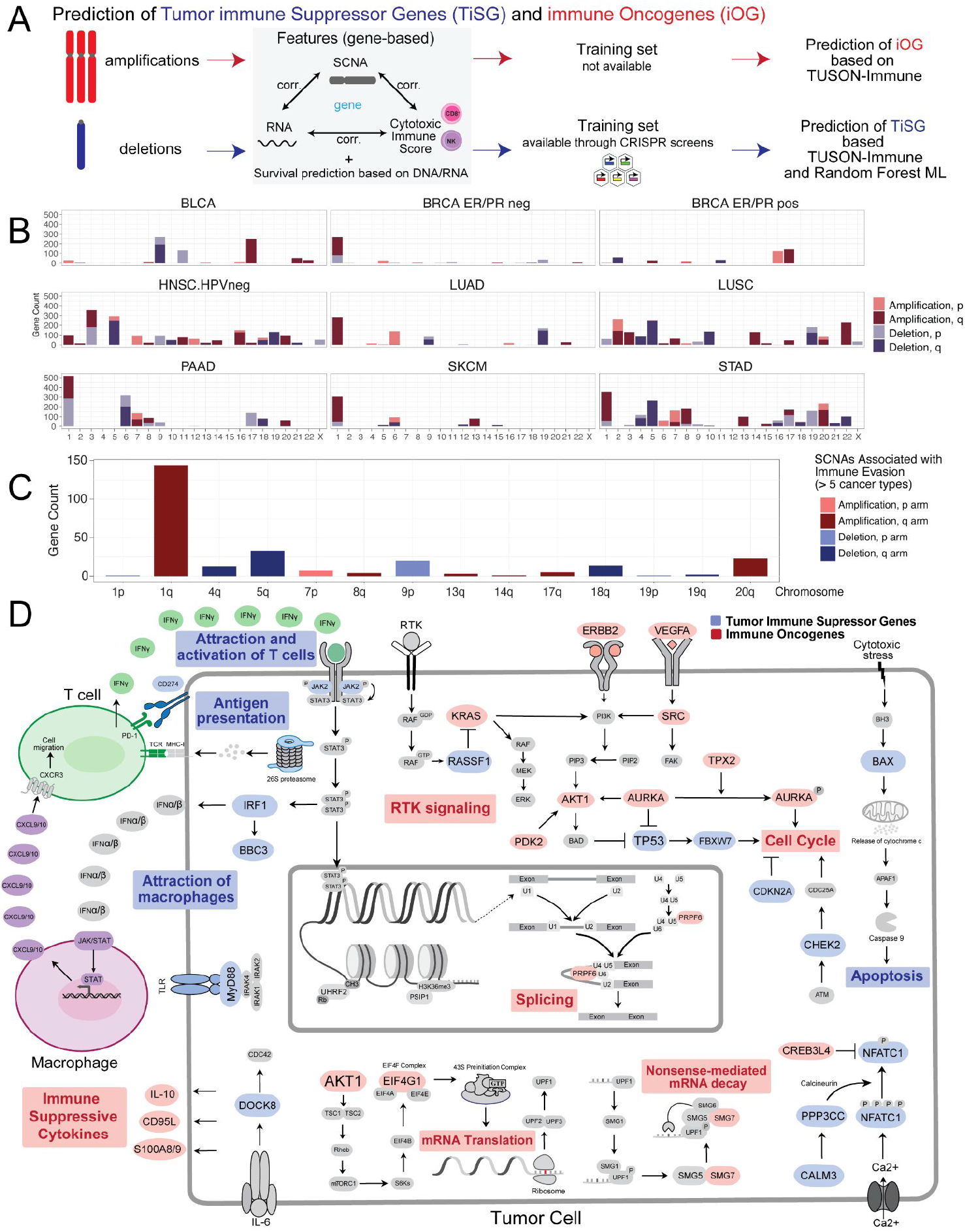
TUSON-Immune prediction of Tumor immune Suppressor Genes (TiSG) and immune Oncogenes (iOG). (A) Schematic of the TUSON-Immune framework: gene-level features (DNA– RNA, RNA–IS, DNA–IS correlations) are used to predict iOGs from amplifications and TiSGs from deletions; TiSG prediction is further refined using a Random Forest classifier trained on CRISPR screen–validated genes. (B) Bar charts showing the chromosomal distribution of predicted TiSGs and iOGs per arm for selected tumor types. (C) Genes predicted as TiSG or iOG in more than five cancer types, grouped by chromosomal arm. (D) Pathway diagram illustrating key TiSG and iOG pathways in the context of tumor–immune interactions. Blue nodes represent TiSGs; red nodes represent iOGs.

Chromosome-level distributions of predicted TiSGs and iOGs are shown for a selection of tumor types (Figure 4B). Across tumor types, chromosome 9p harbored the highest number of TiSGs (Figure 4B–C), consistent with the prominent immune-cold associations of 9p loss. Chromosome 1q had the highest number of predicted iOGs across tumor types, as visible in Figure 4C where 1q shows the tallest amplification bar among arms significant in five or more cancer types. Melanoma (SKCM) is among the tumor types included in Figure 4B.

Pathway analysis of genes predicted as TiSGs or iOGs in three or more tumor types revealed distinct biological enrichments (Figure 4D). TiSGs were enriched for genes involved in cytotoxic T-cell and NK-cell attraction and activation (IFNγ signaling, JAK/STAT pathway, antigen presentation via the proteasome), macrophage recruitment (CXCL10), and pro-apoptotic signaling (BAX, BBC3). Key TiSG candidates on 9p include *JAK2*, interferon genes, and *IRF1*. In contrast, iOGs were enriched for immune-suppressive cytokine secretion (IL-10, S100A8/9), RTK signaling (KRAS, AKT1), cell-cycle promotion (AURKA, ERBB2), and RNA processing pathways (splicing, nonsense-mediated decay, translation initiation). On chromosome 1q, candidate iOGs include *S100A8*/*S100A9* (1q21), which promote the recruitment of immunosuppressive myeloid cells.

Together, these results suggest a mechanistic model in which 9p loss depletes a cluster of immune-activating TiSGs (IFN pathway, JAK2, antigen presentation), directly impairing cytotoxic immune cell recruitment, while 1q gain amplifies iOGs such as S100A8/9 that suppress anti-tumor myeloid and lymphoid responses.

### Chromosome 1q gain predicts poor survival after immunotherapy in melanoma across two independent cohorts

To determine whether immune-cold SCNA events translate into clinical resistance to ICB in melanoma, we performed survival analysis in patients treated with anti-PD-1/PD-L1 therapy. We first generated a genome-wide map of SCNA–survival associations in melanoma using the MSK-IMPACT cohort, applying Cox proportional hazard models to patients stratified by arm-level SCNA status (Figure 5A). The signed −log_10_(p) landscape showed that chromosome 1q gain was the most prominent event associated with worse survival, with the peak signal concentrated on the 1q arm. Other events including chromosome 6p gain and several losses were also associated with altered survival, consistent with the immune-cold associations identified in the KaryoTME analysis.

**Figure 5.**
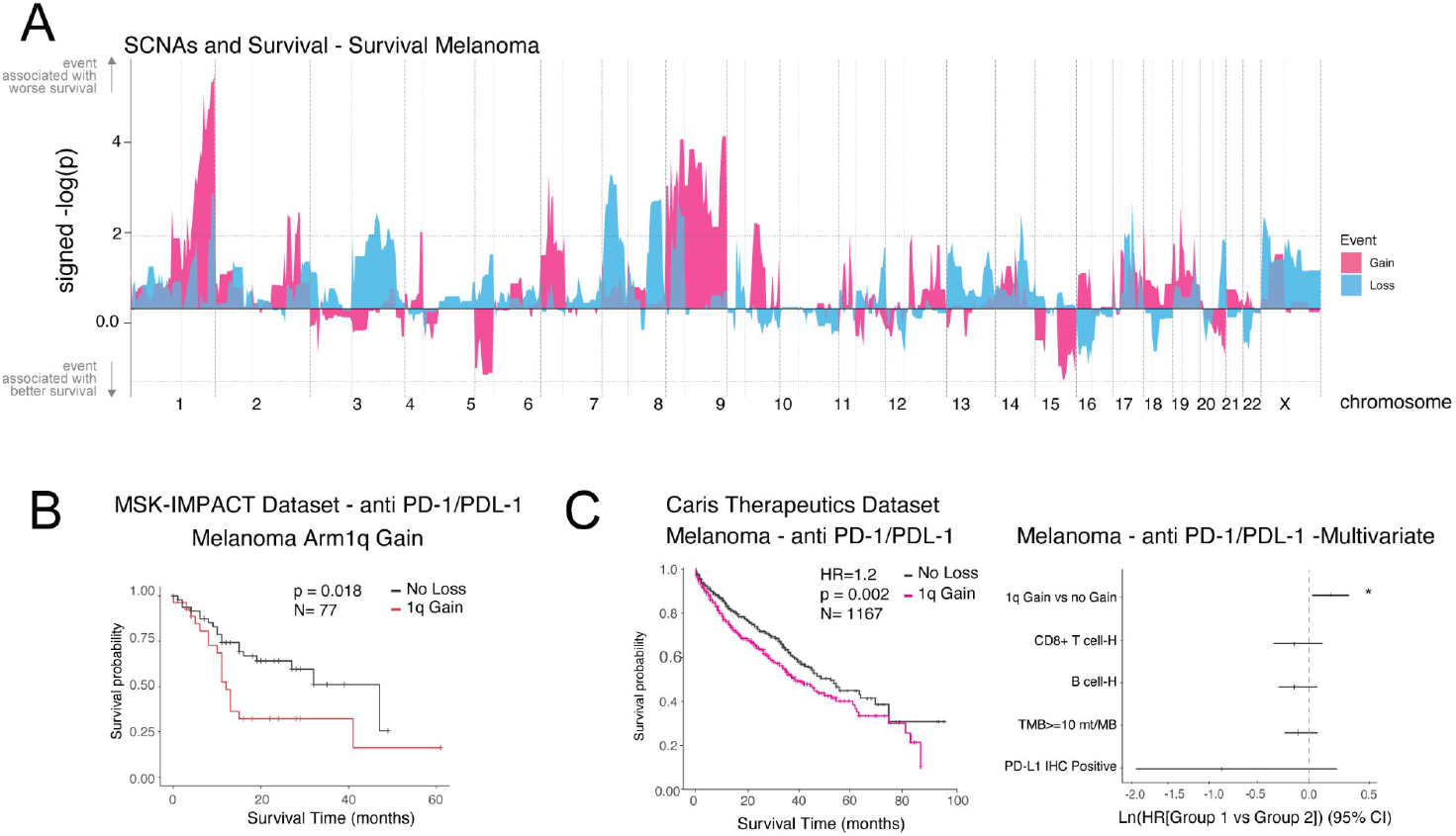
Chromosome 1q gain predicts poor survival after immunotherapy in melanoma. (A) Genome-wide SCNA–survival association landscape in melanoma (MSK-IMPACT cohort). Y-axis: signed −log_10_(p) from Cox regression; events above the midline (positive) are associated with worse survival. Pink: gains; blue: losses. (B) Kaplan–Meier survival curves for melanoma patients treated with anti-PD-1/PD-L1 therapy stratified by arm-level 1q gain status in the MSK-IMPACT cohort (p = 0.018; N = 77). (C) Left: Kaplan–Meier curves for melanoma patients in the Caris Life Sciences real-world cohort treated with anti-PD-1/PD-L1 (HR = 1.2, p = 0.002; N = 1,167). Right: Forest plot from multivariate Cox regression including 1q gain, CD8^+^ T-cell infiltration, B-cell infiltration, TMB ≥10 mut/Mb, and PD-L1 IHC status. Asterisk (*) marks significant association; dashed line at zero (no effect).

Focusing on 1q gain, Kaplan–Meier analysis in the MSK-IMPACT melanoma cohort (N = 77 ICB-treated patients) demonstrated significantly shorter survival for patients with 1q gain compared to those without (p = 0.018; Figure 5B). We replicated this finding in an independent, much larger real-world cohort from Caris Life Sciences, which included 1,167 melanoma patients treated with anti-PD-1/PD-L1 therapy (Figure 5C). In this cohort, 1q gain was significantly associated with worse survival (HR = 1.2, p = 0.002).

To assess whether the prognostic impact of 1q gain is independent of established clinical and immunological biomarkers, we performed multivariate Cox regression including five covariates: 1q gain status, CD8^+^ T-cell infiltration (high vs. low), B-cell infiltration, tumor mutational burden (TMB ≥10 mut/Mb), and PD-L1 immunohistochemistry status. Strikingly, 1q gain was the only variable significantly associated with poor outcome in this multivariate model (Figure 5C, right panel; asterisk marks significance). The other biomarkers, including CD8^+^ T-cell abundance and PD-L1 status, did not reach significance in this analysis, consistent with the well-documented limitations of these markers in melanoma. These results establish chromosome 1q gain as a robust, independent predictor of ICB resistance in melanoma.

## Discussion

In this study, we applied the KaryoTME framework to systematically map the relationship between arm- and focal-level SCNAs and the tumor immune microenvironment, with a focus on melanoma. Our main findings are: (1) chromosome 1q gain and 9p loss are the top immune-cold SCNA events in melanoma; (2) 9p loss primarily depletes cytotoxic T and NK cells while 1q gain depletes both CD8^+^ T cells and NK cells in melanoma and shows broad associations with macrophage depletion pan-cancer; (3) TUSON-Immune identifies S100A8/S100A9, JAK2, and IFN pathway genes as key candidate drivers; and (4) 1q gain is an independent predictor of poor survival after ICB therapy in two large melanoma cohorts.

The prominence of 9p loss as an immune-suppressive SCNA in melanoma is consistent with prior findings linking this alteration to reduced T-cell infiltration and ICB resistance in melanoma and other solid tumors. Our analysis adds resolution by showing that 9p loss preferentially depletes NK and CD8^+^ T cells relative to myeloid populations, and that the cytobands 9p21 (interferon genes, *CDKN2A*), 9p23, and 9p24 (JAK2, PD-L1/PD-L2) all contribute to this phenotype. Critically, the immunosuppressive effect appears to arise from the cumulative loss of multiple immune-activating genes along the arm, rather than any single gene, consistent with a multi-gene synergy model.

The discovery that chromosome 1q gain is a robust, independent predictor of ICB resistance in melanoma is clinically significant. Chromosome 1q is one of the most frequently gained chromosomal arms in human cancer and carries a dense cluster of genes implicated in oncogenesis and immune suppression. Our TUSON-Immune analysis prioritizes S100A8 and S100A9 (1q21) as candidate iOGs, consistent with their roles in promoting myeloid-derived suppressor cell recruitment and polarizing macrophages toward an immunosuppressive phenotype. Other candidate iOGs on 1q include genes involved in cell-cycle regulation and oncogenic signaling. Importantly, the association of 1q gain with poor ICB outcomes was replicated across two independent cohorts spanning nearly 1,250 melanoma patients and remained significant after adjusting for CD8^+^ T cells, B cells, TMB, and PD-L1 (all standard clinical biomarkers currently used to guide ICB decisions). This suggests that 1q gain captures a distinct biological axis of resistance not reflected in these existing markers.

Our data also raise interesting questions about the mechanisms by which 1q gain promotes immune evasion. The observed association with reduced NK and CD8^+^ T-cell proportions in the scRNA-seq validation dataset, combined with the broad SCNA-macrophage associations in the pan-cancer analysis, suggests a model in which 1q gain both directly impairs cytotoxic cell recruitment and remodels the myeloid compartment toward an immunosuppressive state. Whether these effects are mediated predominantly through the secreted proteins S100A8/A9, through other 1q-encoded oncogenic regulators, or through combined dosage effects of many genes simultaneously will require functional investigation.

Taken together, our findings suggest that routine assessment of chromosome 1q copy number status may improve patient stratification for ICB therapy in melanoma, either independently or in combination with existing biomarkers. As whole-genome or targeted sequencing is increasingly performed as standard of care in advanced melanoma, incorporating arm-level SCNA data into biomarker panels is now practically feasible. Future prospective studies should validate the predictive value of 1q gain in the context of current first-line combination regimens, and functional studies should clarify which genes on 1q are the principal drivers of immune evasion.

## Methods

### Patient datasets and data sources

Clinical, copy number, and gene expression data from The Cancer Genome Atlas (TCGA) were downloaded from The Broad GDAC Firehose (https://gdac.broadinstitute.org/). Single-cell RNA-seq data from skin melanoma were obtained from Biermann et al. (2022). Clinical and SCNA data from patients treated with immune checkpoint blockade were obtained from the MSK-IMPACT cohort. Real-world clinical and transcriptomic data from melanoma patients were obtained from Caris Life Sciences as described in the text.

### Cytotoxic Immune Score

The cytotoxic Immune Score (IS) was computed from TCGA RNA-seq data as a ranked-sum score based on the expression of seven cytotoxic genes: CD247, CD2, CD3E, GZMH, NKG7, PRF1, and GZMK. Each gene was ranked across all samples within a dataset; ranks were summed per sample and then re-ranked to obtain the final IS. Samples in the bottom 30th percentile were classified as immune cold; samples in the top 30th percentile as immune hot.

### SCNA calling and aneuploidy score

Raw copy number calls from GISTIC2 were adjusted for tumor purity and ploidy using ABSOLUTE algorithm estimates. A chromosomal segment was classified as a gain if log^2^(CN) > 0.2, and as a loss if log^2^(CN) < −0.2. The aneuploidy score (AS) was computed as the sum of absolute log^2^(CN) values across all arms, cytobands, or genes (matching the level of analysis).

### Logistic regression models

Logistic regression models were run independently per tumor type. Models predicted immune score (IS, binarized as above) from arm-, cytoband-, or gene-level log^2^(CN) values, with the AS included as a covariate. Gain and loss models were run separately: samples with losses were excluded from gain models, and samples with gains were excluded from loss models. Four models were run per tumor type per genomic level: two predicting immune cold (one for gains, one for losses) and two predicting immune hot. P-values were adjusted using the Benjamini–Hochberg (BH) method within each tumor type and SCNA type independently.

### xCell deconvolution and cell-type regression models

Immune and stromal cell-type proportions were estimated from TCGA bulk RNA-seq data using xCell. Linear regression models were then run for each cell type using the same SCNA covariates and aneuploidy score framework as the IS models. CNV–cell type associations with p > 0.2 or directionally inconsistent gain/loss effects were excluded.

### Single-cell RNA-seq analysis in melanoma

We analyzed published single-cell RNA-seq data from Biermann et al. (2022) profiling the melanoma TME (21 patients). Cell type annotations were obtained from the published metadata. To infer large-scale copy number alterations at the single-cell level, inferCNV was applied using stromal and non-malignant cells as reference. Immune cell proportions were calculated per patient. Statistical differences in immune cell proportions between SCNA status groups were assessed by t-tests.

### TUSON-Immune prediction of TiSGs and iOGs

Each gene was evaluated using three LASSO-selected parameters: DNA:IS correlation, RNA:IS correlation, and DNA:RNA correlation. For iOG prediction, genes in amplified regions were required to show adjusted p < 0.05 and positive rho for all three parameters. For TiSG prediction, genes in deleted regions were required to show adjusted p < 0.05 with positive DNA:RNA rho and negative DNA:IS and RNA:IS rho. A Random Forest classifier trained on a curated TiSG gold standard (Li et al. 2022, 65 genes from 17 CRISPR screens) was used as a complementary TiSG predictor, achieving AUROC of 0.87 (cross-validation) and 0.89 (test set).

### Survival analysis

Survival analysis in the MSK-IMPACT cohort was performed using Cox proportional hazard models for ICB-treated melanoma patients. SCNAs were categorized as Gain (> 0.3), Loss (< −0.3), or Neutral. Kaplan–Meier curves were generated with log-rank tests. In the Caris Life Sciences real-world cohort, overall survival was derived from real-world evidence data. Chromosome 1q copy number was inferred from whole-transcriptome sequencing using a panel of 50 genes with high DNA–RNA concordance in TCGA melanoma. Patients in the top quartile of 1q expression score were classified as 1q-gain. Multivariate Cox regression included CD8^+^ T-cell fraction, B-cell fraction, PD-L1 IHC status, and TMB (≥10 mut/Mb) as additional covariates. All p-values were adjusted by the false discovery rate.

## Acknowledgements

We thank members of the Davoli laboratory for their discussions and support, and Spencer Bruce for help assembling Figure 1. We thank Trinity de Leon for expert help in figure editing. This work was supported by grants from the National Institutes of Health (R37CA248631, R01HG012590, and R01DK135089), the Pershing Square Sohn Prize for Young Investigators in Cancer Research, the Melanoma Research Alliance Young Investigator Award to T.D., and a collaborative grant from the NFCR.

